# Blocking mitochondrial pyruvate import causes energy wasting via futile lipid cycling in brown fat

**DOI:** 10.1101/841551

**Authors:** Michaela Veliova, Caroline M. Ferreira, Ilan Y. Benador, Anthony E. Jones, Brandon R. Desousa, Kiana Mahdaviani, Rebeca Acín-Pérez, Anton Petcherski, Ajit S. Divakaruni, Marc Prentki, Barbara E. Corkey, Marc Liesa, Marcus F. Oliveira, Orian S. Shirihai

**Affiliations:** Department of Molecular and Medical Pharmacology, David Geffen School of Medicine at UCLA, Los Angeles, CA, USA; Department of Medicine, Division of Endocrinology, David Geffen School of Medicine at UCLA, Los Angeles, CA, USA; Instituto de Bioquímica Médica Leopoldo de Meis, Universidade Federal do Rio de Janeiro, Rio de Janeiro, RJ, Brazil; Nutrition and Metabolism, Graduate Medical Sciences, Boston University School of Medicine, Boston, MA, USA; Department of Nutrition, and Department of Biochemistry and Molecular Medicine, University of Montreal, QC, Canada

## Abstract

Futile lipid cycling is an ATP-wasting process proposed to participate in energy expenditure of mature fat-storing white adipocytes, given their inability to oxidize fat. The hallmark of activated brown adipocytes is to increase fat oxidation by uncoupling respiration from ATP synthesis. Whether ATP-consuming lipid cycling can contribute to BAT energy expenditure has been largely unexplored. Here we find that pharmacological inhibition of the mitochondrial pyruvate carrier (MPC) in brown adipocytes is sufficient to increase ATP-synthesis fueled by fatty acid oxidation, even in the absence of adrenergic stimulation. We find that elevated ATP-demand induced by MPC inhibition results from activation of futile lipid cycling. Furthermore, we identify that glutamine consumption and the **M**alate-**A**spartate **Sh**uttle are required for the increase in **E**nergy **E**xpenditure induced by MPC inhibition in **B**rown **A**dipocytes (MAShEEBA). These data demonstrate that futile energy expenditure through lipid cycling can be activated in BAT by altering fuel availability to mitochondria. Therefore, we identify a new mechanism to increase fat oxidation and energy expenditure in BAT that bypasses the need for adrenergic stimulation of mitochondrial uncoupling.

## 1. INTRODUCTION

Brown adipose tissue (BAT) is the main site of inducible thermogenesis. [1–4]. Fat oxidation is activated in BAT when the sympathetic nervous system releases norepinephrine after sensing cold temperatures, which activates adrenergic receptors on BAT to increase thermogenesis. This activation causes a stimulation of lipolysis, mediated by Protein Kinase A actions on lipolytic proteins, ultimately activating mitochondrial uncoupling protein 1 (UCP1) by free fatty acids [1,5]. UCP1 is exclusively expressed in thermogenic adipocytes and dissipates the mitochondrial proton gradient to generate heat, instead of ATP [1]. Numerous studies have shown the beneficial effects of BAT activation in the treatment of metabolic disorders [6]. Although recent publications have proposed mechanisms to increase energy expenditure in BAT independent of hormonal stimulation [7–10], currently there is only one known pharmacological approach to increase BAT energy expenditure in humans. This one approach involves adrenergic stimulation, thus being an approach that activates other tissues and provokes substantial side effects.

Increased glucose uptake is a hallmark of BAT activation, as shown by ^18^flurodeoxyglucose positron emission tomography–computed tomography (^18^F-PET-CT) images in humans and mice [2–4,11]. The end-product of glycolysis, pyruvate, has multiple metabolic fates in the cytosol and in the mitochondria. Mitochondrial pyruvate metabolism requires its transport to the mitochondrial matrix, which is accomplished via the mitochondrial pyruvate carrier (MPC). The MPC, a protein complex located in the inner mitochondrial membrane, enables pyruvate transport to the mitochondrial matrix [12,13]. Several studies have shown that MPC mediates metabolic re-wiring and determines fuel preferences for mitochondrial oxidation with direct consequences to cell function and fates [14–18]. A major mechanism is that pyruvate oxidation allows citrate synthesis, the precursor of malonyl-CoA, which is required for *de novo* lipogenesis and inhibits long chain fatty acid oxidation via its action on carnitine palmitoyl transferase 1 (CPT1). The role of mitochondrial pyruvate oxidation driven by MPC activity in brown adipocytes remains elusive, as the assumptions are that; i) pyruvate might not be required as an oxidative fuel in BAT mitochondria, given that activated brown adipocytes mostly oxidize fatty acids and ii) the preferred fate of pyruvate from increased glycolysis in BAT would be lactate, to allow glycolysis to compensate and provide the ATP not produced by uncoupled mitochondria. To further test these assumptions, we determined the effects of blocking MPC in brown adipose energy expenditure. Here, we show that pharmacological and genetic inhibition of the MPC activates coupled fat oxidation, thus demonstrating that MPC inhibition increases ATP demand and fat expenditure. The increased flux in coupled fatty acid oxidation induced by MPC blockage is supported by glutamine consumption and the malate-aspartate shuttle (MASh), a previously unstudied metabolic pathway in BAT. Interestingly, we show that acute MPC blockage has an additive effect promoting energy expenditure in activated brown adipocytes, suggesting that MPC inhibition enhances fatty acid oxidation for ATP synthesis in mitochondria in which UCP1 was not activated. Finally, we demonstrate that ATP-demanding glycerolipid/ free fatty acid (GL/FFA) cycling is responsible for increased ATP demand in response to MPC inhibition. Thus, we show that MPC activity might be limiting BAT energy expenditure both in resting and activated brown adipocytes, rather than facilitating it. We conclude that MPC activity regulates BAT energy metabolism and its inhibition may represent a novel target to increase fat oxidation and energy expenditure with potential benefits for metabolic diseases.

## 2. RESULTS

### 2.1. Pharmacological inhibition of the MPC in non-stimulated brown adipocytes increases fatty acid oxidation and energy expenditure

To determine the role of the mitochondrial pyruvate import in BAT fuel preference, we sought to assess the effects of MPC inhibition on fatty acid utilization in non-stimulated brown adipocytes. Mitochondrial pyruvate import was blocked using a pharmacological inhibitor that covalently binds the MPC, UK5099 [19]. UK5099 was used at a concentration range demonstrated to have specific effects on mitochondrial pyruvate import [19]. To test fatty acid consumption, we provided the fluorescent fatty acid analog Bodipy C12 558/568 (Bodipy C12) to primary brown adipocytes, which can be oxidized and/or incorporated into lipid droplets (LDs) as triacylglyceride (TAGs). Intracellular lipids of primary brown adipocytes were stained overnight with Bodipy C12 and labeled fatty acid trafficking was measured upon 120 min of UK5099 treatment. Live cell imaging shows that lipid droplet area was significantly reduced upon UK5099 treatment, suggesting increased lipolysis (Figure 1A). Reduced lipid droplet content by MPC inhibition followed the same trend as for NE-induced activation of lipolysis in brown adipocytes (Figure 1A). Next, we tested whether MPC inhibition increased utilization of extracellular fatty acids by measuring the fate of exogenous fluorescently labeled fatty acid using thin layer chromatography (TLC). We pulsed brown adipocytes with Bodipy-C12 for 24 h in the presence of 100 nM UK5099, 1 µM norepinephrine (NE) or vehicle. Lipids were extracted from both media and cells and resolved by TLC (Figure 1B) [20,21]. Treatment with UK5099 caused an increase in Bodipy C12 uptake from the media similarly to NE, indicating that brown adipocytes increase fat utilization upon MPC inhibition from both intracellular- and extracellular source (Figure 1B-C). We then quantified the amount of TAGs synthesized per free C12 taken up from the media and found that both UK5099 and NE reduced fatty acid esterification to TAGs (Figure 1C). Altogether, these data provide evidence that MPC inhibition promotes lipid uptake by brown adipocytes and a switch towards fatty oxidation rather than storage as TAG.

**Figure 1:**
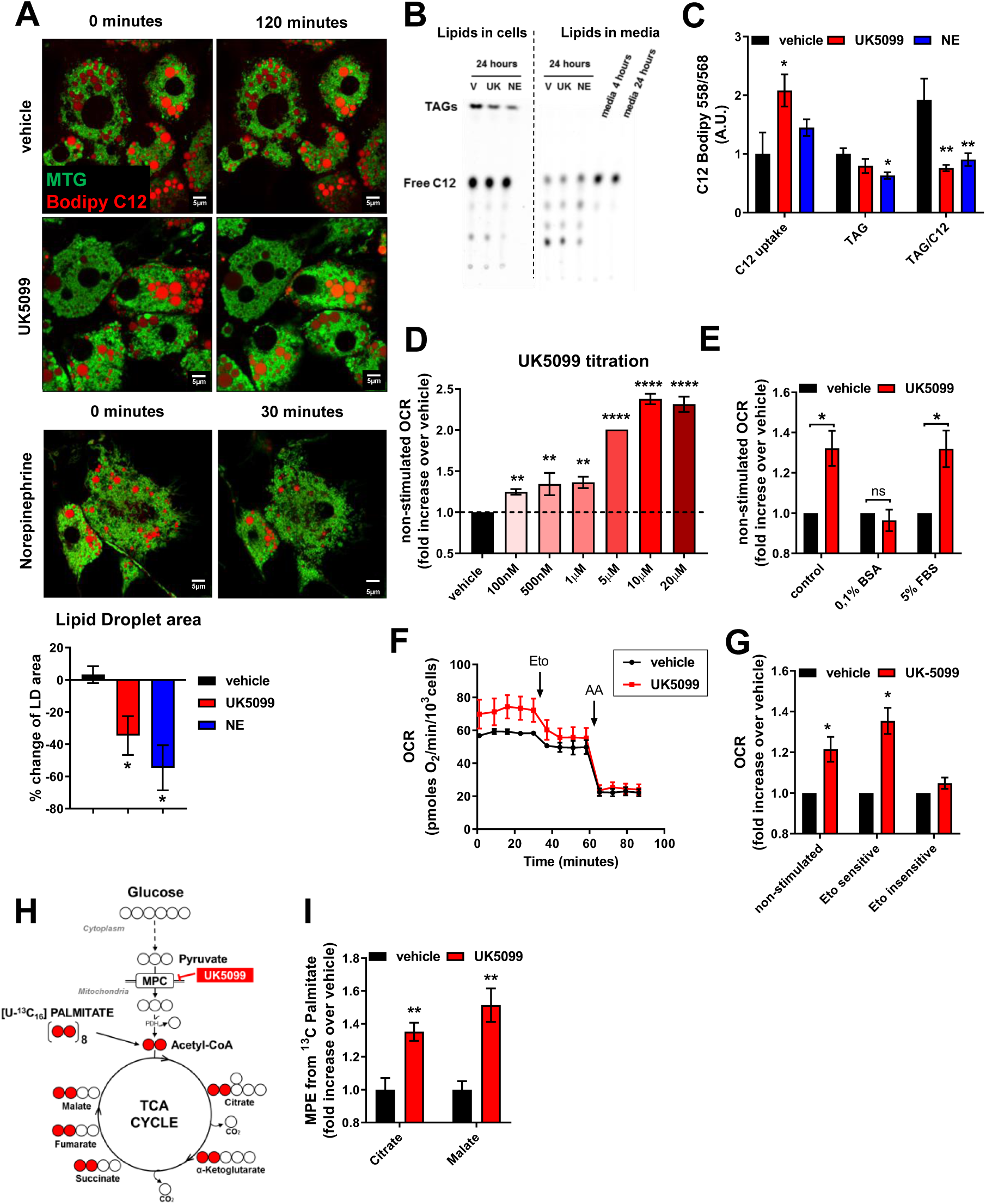
Pharmacological inhibition of the MPC in non-stimulated brown adipocytes increases energy expenditure fueled by fatty acids. **(A)** Live-cell super-resolution confocal imaging of primary brown adipocytes pre-stained overnight with Bodipy C12 558/568. Cells were stained with mitotracker green (MTG) prior to imaging. Quantification of changes in lipid droplet cross-sectional area with indicated treatments. Data are represented as percentage change in area at time=120 min compared to time=0. Data represent n=20-30 cells from 3 individual experiments. * p < 0.05 by ANOVA. **(B)** Representative thin-layer chromatography (TLC) plate of lipids extracted from extracellular content (cell culture media) and primary brown adipocytes treated with vehicle (V), 100 nM UK5099 (UK) or 1 µM norepinephrine (NE). Primary brown adipocytes were incubated with BODIPY C12 558/568 (C12) in presence of vehicle (V), UK5099 (UK) or norepinephrine (NE) for 24 h to assess triacylglyceride (TAG) synthesis (Lipids in cells) or fatty acid uptake (Lipids in media). The relative polarity of the lipid species determines the motility, with TAG migrating the highest. **(C)** Quantification of TAG per amount of free fatty acid uptake from n=4 individual experiments. Note that UK5099 decreases the amount of TAG build per free fatty acid similarly to NE. Data were normalized to vehicle for each individual experiment. ns p>0.05, ** p < 0.01 by ANOVA. **(D)** Fully differentiated primary brown adipocytes were pre-treated with vehicle (DMSO) UK-5099 at indicated concentrations for 2 h. Oxygen consumption rates (OCR) were measured in Seahorse base media supplemented with 5 mM glucose and 3 mM glutamine in the presence of vehicle or UK-5099. Quantification of non-stimulated OCR. Data were normalized to vehicle for each individual experiment. ** p < 0.01, **** p < 0.0001 compared to vehicle by ANOVA. **(E)** Oxygen consumption rates (OCR) were measured in Seahorse base media supplemented with 5 mM glucose and 3 mM glutamine in the presence of vehicle or UK-5099. Quantification of non-stimulated OCR of brown adipocytes from n=3-7 experiments. OCR were measured in Seahorse base media with glucose and glutamine (control), supplemented with 0.1% fatty acid free BSA or supplemented with 5% FBS. Data were normalized to vehicle for each individual experiment. ns p>0.05, * p < 0.05 compared to vehicle by Student’s t test. **(F)** Brown adipocytes were treated with 100 nM UK5099 or vehicle. 40 µM etomoxir (Eto) and 4 µM antimycin a (AA) when indicated. Data shows representative OCR traces from n=6 technical replicates. **(G)** Quantification of etomoxir sensitive, and etomoxir insensitive OCR from n=4 experiments. Data were normalized to vehicle for each individual experiment. * p < 0.05 compared to vehicle by Student t test. **(H)** Schematic representation of metabolite tracing using [U-^13^C_16_] palmitate. **(I)** [U-^13^C_16_] palmitate tracing in fully differentiated primary brown adipocytes treated with 5 µM UK5099 or vehicle for 24 h. Data shows mole percent enrichment (MPE) of isotope labeled substrate into respective metabolite. Data were normalized to vehicle for each individual replicate and represent n=6 technical replicates from 2 individual experiments.

Given that the data suggest that MPC inhibition results in increased lipolysis and uptake of extracellular fatty acids, and thus higher fatty acid availability to mitochondria, we reasoned that blocking mitochondrial pyruvate import might mimic the effects of NE and increase energy expenditure by activating mitochondrial respiration. Thus, we measured the effects of MPC inhibition on oxygen consumption rates (OCR) in non-stimulated brown adipocytes. Remarkably, inhibition of the MPC resulted in a dose-dependent increase in OCR, with maximal effects observed at 10 µM UK5099 (Figure 1D). This observation raised the question as to which substrate is fueling the observed increase in mitochondrial respiration. Given that both lipolysis and fatty acid uptake were increased under UK5099 treatment (Figure 1A-C), we tested the requirement for fatty acids in UK5099-induced increase in OCR. We first reasoned that, if increased energy expenditure caused by UK5099 requires a rise in intracellular levels of fatty acids, then their sequestration would prevent the boost on respiratory rates upon MPC inhibition. To test this prediction, we supplemented the respirometry media with 0.1 % BSA-FAF (fatty acid free), which would remove the excess intracellular free fatty acids [22]. Figure 1E shows that supplementation with BSA-FAF reversed the stimulatory effects of UK5099 on basal respiration, when compared to control media. However, when respirometry media was supplemented with 5 % FBS, which matches similar albumin concentrations as in the FAF BSA-treated group but contains fatty acids, the effects of UK5099 increasing basal respiration were restored (Figure 1E). These data suggest that fatty acid oxidation supports increased OCR when the MPC is inhibited.

To further test the hypothesis that UK5099 increases OCR by promoting β-oxidation with a more direct approach, we used etomoxir to block the rate limiting step of ß-oxidation catalyzed by carnitine palmitoyl transferase 1 (CPT1) [23]. The contribution of β-oxidation to OCR was then assessed by quantifying the effects of etomoxir on respiratory rates before and after its injection in Seahorse cartridges (named hereafter as the “etomoxir-sensitive OCR”). Etomoxir was used at 40 μM, a concentration that we have determined to inhibit CPT1 dependent OCR with only minimal effects on other mitochondrial oxidative pathways (Figure S1). Remarkably, the etomoxir sensitive component of the OCR of cells treated with UK5099 was significantly higher as compared to vehicle-treated cells, strengthening our assertion that MPC inhibition promotes fatty acid oxidation (Figures 1F-G). We observed that the OCR component insensitive to etomoxir, which reflects the metabolic pathways other than β-oxidation that contributes to respiration, were not affected by MPC inhibition. This further confirms the selective nature of the metabolic shift in brown adipocytes towards fatty acid oxidation induced by MPC inhibition.

Since the results using the Seahorse bioanalyzer and measuring etomoxir-sensitivity only reflect fuel dependence but do not assert whether MPC inhibition leads to a fuel preference switch towards fatty acid oxidation in brown adipocytes, we analyzed the incorporation of palmitate-derived carbons into TCA cycle intermediates measured by gas-chromatography/mass spectrometry (GC/MS) (Figure 1H). [U-^13^C_16_] Palmitate was provided to fully differentiated primary brown adipocytes for 24 h, together with 5 µM UK5099 or vehicle. Polar metabolites were extracted and the enrichment of ^13^C in TCA cycle metabolites was quantified. Figure 1I shows that UK5099 increased incorporation of ^13^C from labelled palmitate into TCA cycle metabolites citrate and malate, indicating that inhibition of the MPC increases, under non-stimulated conditions, the utilization of fatty acids and the production of acetyl-CoA that enters the TCA cycle. Thus, our data indicate that pharmacological inhibition of the MPC increases energy expenditure in brown adipocytes by increasing fatty acid oxidation, in the absence of NE-stimulation.

### 2.2. Stimulation of energy expenditure by norepinephrine is enhanced by acute MPC inhibition

Given that there is an increase in OCR upon MPC inhibition under non-stimulated conditions, we tested the effects of UK5099 on norepinephrine-stimulated respiration in brown adipocytes. OCR was measured before (non-stimulated) and after exposure to 1 µM norepinephrine (NE-stimulated). Treatment of brown adipocytes with 100 nM UK5099 increased NE-stimulated OCR (Figures 2 A-B). To determine the relative contribution of fatty acid oxidation to NE-stimulated energy expenditure, we tested the effect of etomoxir on OCR after NE stimulation. To more specifically assess the contribution of fatty acid oxidation to uncoupled respiration we inhibited ATP synthase by injecting oligomycin before etomoxir. Together this protocol allows determination of the effect of MPC inhibition on uncoupled respiration energized by fatty acids in NE-stimulated brown adipocytes. The component of NE-stimulated OCR sensitive to etomoxir was significantly higher in cells treated with UK5099 compared to vehicle-treated cells, indicating that even in NE-stimulated cells UK5099 further promotes an increase in fatty acid oxidation (Figure 2A-B). To confirm that the UK5099 effects were caused by their action on MPC, we tested the effects of reducing the expression of the MPC on energy expenditure and fatty acid oxidation. The MPC is a protein heterodimer, comprised the subunits MPC1 and MPC2. The expression of both subunits is required for functional pyruvate import, and deletion of either of the subunits was shown to lead to impairment of mitochondrial pyruvate import [12,13,24]. Therefore, brown adipocytes were transfected with an siRNA for MPC1 (MPC1-KD) or a scramble RNA (Scramble), and successful knock-down of MPC1 was confirmed by Western blot (Figures 2C-D) and by qPCR (Figure 2E). Both MPC1 and MPC2 protein levels were reduced when cells were transfected with MPC1 siRNA compared to scramble RNA (Figure 2D). However only mRNA levels of MPC1 but not MPC2 were reduced in cells transfected with MPC1 siRNA (Figure 2E). Next, we tested the effects of MPC1-KD on energy expenditure and fatty acid oxidation. Assessment of OCR in non-stimulated brown adipocytes showed that MPC1-KD increased energy expenditure compared to Scramble RNA, thereby confirming the observed effect of pharmacological MPC inhibition (Figures 2F-G). Similar to treatment with UK5099, MPC1-KD increased the proportion of energy expenditure dependent on fatty acid oxidation (Figure 2G). Interestingly, MPC1-KD did not change NE-stimulated OCR compared to Scramble (Figure 2G). The apparent discrepancy between pharmacological and genetic interference might suggest that short-term versus long-term reduction of MPC activity cause different metabolic phenotypes, as the siRNA transfection represents are more chronic MPC inhibition, as opposed to UK5099 treatment which is a 2 h acute inhibition of pyruvate import. To further confirm that the effects of UK5099 on brown adipocyte energy expenditure are on-target, we treated Scramble RNA and MPC1-KD cells with 100 nM UK5099. Cells transfected with scramble RNA had increased basal OCR following UK5099 treatment, which agrees with the results using non-transfected cells (Figure 2H). However, the stimulatory effect of UK5099 on OCR was lost in MPC1-KD cells (Figure 2H), indicating that increased energy expenditure caused by UK5099 treatment is dependent on MPC.

**Figure 2:**
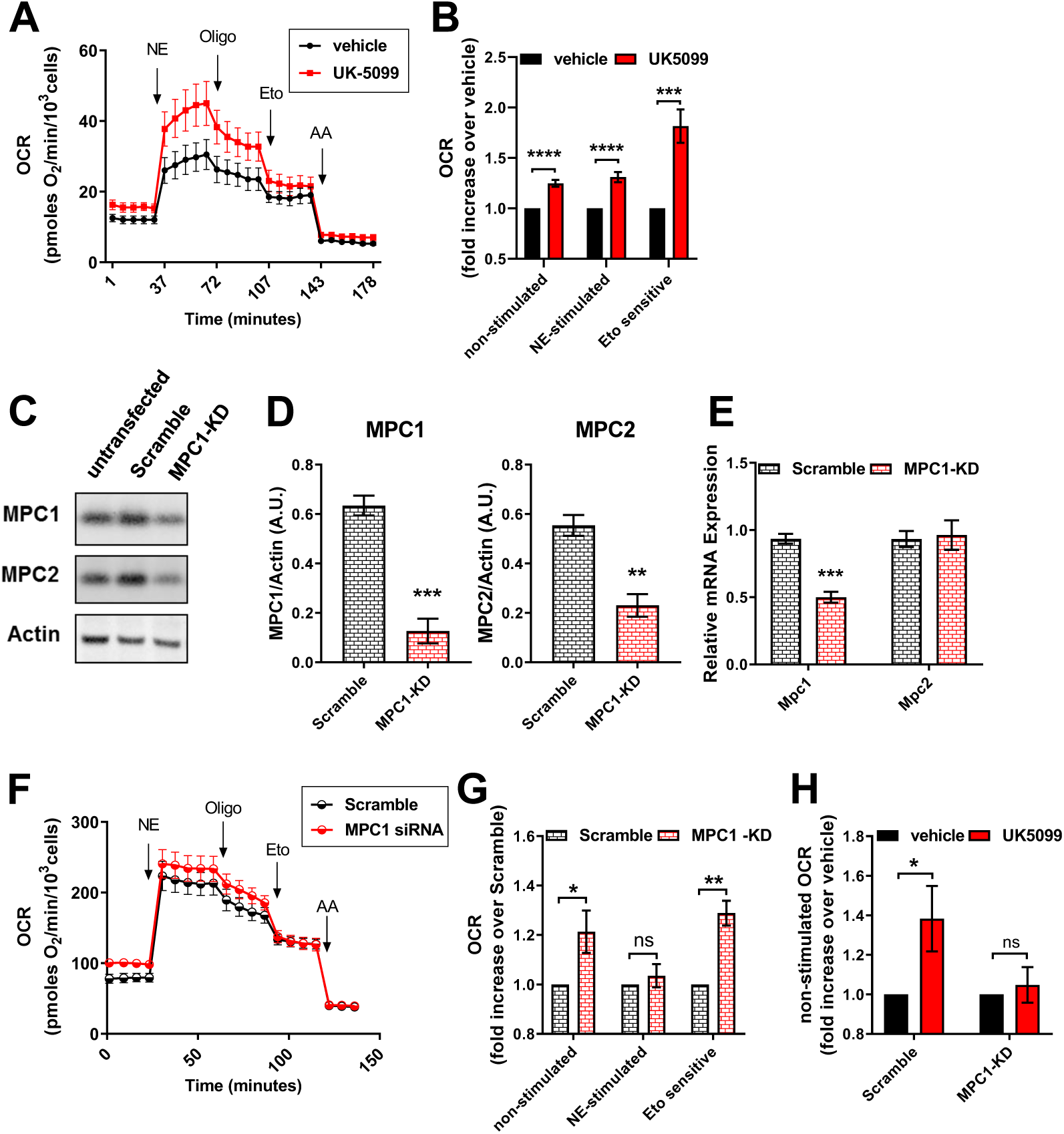
Stimulation of energy expenditure by norepinephrine is enhanced by acute MPC inhibition. **(A)** Fully differentiated primary brown adipocytes were pre-treated with vehicle (DMSO) or 100 nM UK5099 for 2 h. OCR were measured in Seahorse base media supplemented with 5 mM glucose and 3 mM glutamine in the presence of vehicle or UK-5099. 1 µM norepinephrine (NE), 4 µM oligomycin A (Oligo), 40 µM etomoxir (Eto) and 4 µM antimycin a (AA) were injected when indicated. Data shows representative OCR traces from n=4 technical replicates. **(B)** Quantification of non-stimulated, NE-stimulated and Eto-sensitive OCR (n=9 experiments). The effects of 100 nM UK5099 treatment for 2 h were normalized to vehicle for each experiment. *** p < 0.001, **** p < 0.0001 compared to vehicle by Students t test. **(C)** Representative Western Blot for MPC1 and MPC2 of differentiated primary brown adipocytes transfected with MPC1 siRNA (MPC1 KD) or Scramble RNA (Scramble). **(D)** Quantification of MPC1 and MPC2 expression normalized to actin (n=3 experiments). ** p < 0.01 compared to Scramble by Students t test. **(E)** mRNA levels of MPC1 and MPC2 in brown adipocytes transfected with scramble RNA (Scramble) or MPC1 siRNA (MPC1-KD). ** p < 0.01 compared to Scramble by Student’s t-test. **(F)** Representative OCR traces of differentiated primary brown adipocytes transfected with MPC1 siRNA (MPC1 KD) or Scramble RNA (Scramble). OCR were measured in Seahorse base media supplemented with 5 mM glucose and 3 mM glutamine in Scramble RNA or MPC1-KD brown adipocytes. 1 µM norepinephrine (NE), 4 µM oligomycin A (Oligo), 40 µM etomoxir (Eto) and 4 µM antimycin a (AA) were injected when indicated. Data show representative OCR traces from n=3 technical replicates. **(G)** Quantification of basal, NE-stimulated and Eto sensitive OCR (n=6 experiments). Data were normalized to Scramble RNA for each individual experiment. ns p>0.05, * p < 0.05, **** p < 0.0001 compared to Scramble by Student’s t test. **(H)** Quantification of basal OCR in response to UK5099 treatment in Scramble RNA of MPC1 siRNA transfected cells (n=4 individual experiments). Data were normalized to vehicle for each experiment. Note that activating effect of UK5099 on OCR in brown adipocytes is lost by MPC1KD. ns p>0.05, * p < 0.05 compared to vehicle by Student’s t test.

### 2.3. Increased fatty acid oxidation and energy expenditure promoted by MPC inhibition require glutamine metabolism and the activation of malate aspartate shuttle

Fatty acid oxidation generates acetyl-CoA, which is the same product that is derived from pyruvate oxidation in the mitochondria. However, mitochondrial pyruvate metabolism also provides essential carbons to the TCA cycle through CO_2_-fixing reactions such as pyruvate carboxylase, an enzyme located in the mitochondrial matrix that produces oxaloacetate (OAA) for condensation with acetyl CoA and continued TCA cycle flux [25]. Fatty acid-derived carbon enters the TCA cycle only through acetyl CoA and thus cannot directly generate oxaloacetate (Figure 3C). Therefore, oxaloacetate could become limiting when MPC activity is inhibited. Since MPC inhibition did not reduce, but rather increase respiratory rates linked to fatty acid oxidation, we hypothesized that brown adipocytes switch to an alternative source for oxaloacetate when mitochondrial pyruvate transport is limited. Previous work in liver, neurons and myoblasts showed that cells shift towards glutamate metabolism when the MPC is inhibited, to support both biosynthesis and energy demand [14,15,17]. We therefore assessed the dependency of UK5099-stimulated increase in OCR on glutamine in brown adipocytes. UK5099 treatment increased non-stimulated and NE-stimulated OCR only in the presence of glucose and glutamine in combination (Figures 3A-B). When glucose was the sole nutrient source, pharmacological inhibition of the MPC reduced non-stimulated as well as NE-stimulated respiration (Figures 3A-B). These data indicate that glutamine is a necessary nutrient to sustain increased OCR upon MPC inhibition. However, to determine if glutamine is essential for fatty acid oxidation or as an alternative fuel, we determined the contribution of glutamine to fatty acid oxidation by measuring etomoxir-sensitive OCR in the presence and absence of glutamine. Figures 3A and 3B show that increases in etomoxir-sensitive OCR following MPC inhibition only took place when both glucose and glutamine were provided. This indicates that glutamine metabolism is required for the enhancement of fatty acid oxidation upon MPC inhibition. A mechanism by which glutamine provides carbons for the TCA cycle is through the transport of glutamate via the malate aspartate shuttle (MASh) a cyclic pathway that allows the transfer of reduced equivalents to the mitochondrial matrix (Figure 3C). We reasoned that activation of MASh would provide a mechanism to sustain TCA cycle flux, maintain oxaloacetate concentration in a proper range and allow increased fatty acid oxidation to proceed. To determine the role of the MASh in increased oxygen consumption induced by MPC inhibition, we inhibited the glutamic oxalacetic transaminase (GOT) by using aminooxyacetic acid (AOA) [26]. Note that the MASh is dependent on GOT to sustain OAA generation from glutamate, with GOT being commonly blocked to inhibit of MASh activity. UK5099-mediated increase in OCR in non-stimulated brown adipocytes was dose-dependently inhibited by AOA, reaching complete reversal of UK5099-effect at 1 mM AOA (Figure 3D and S3B). In the absence of UK5099, AOA did not have any significant effect on OCR in non-stimulated brown adipocytes (Figure 3D and S3A). To confirm the involvement of MASh in the metabolic effects of MPC inhibition, we silenced the oxoglutarate carrier 1 (SLC25A11, or OGC1), a critical component of the MASh which mediates the exchange of malate and α-ketoglutarate between intermembrane space and mitochondrial matrix (Figure 3C). Adenovirus-mediated knock-down of OGC1 was confirmed by qPCR in primary brown adipocytes (Figure S3C). OGC1 KD prevented the increases in respiratory rates provided by UK5099 treatment under non-stimulated and NE-stimulated conditions, which contrasts with higher respiratory rates in cells transduced with scramble RNA (Figures 3E-F and S3C). To confirm that MASh was required for the activation of fatty acid oxidation under MPC inhibition, we blocked CPT1 activity by using etomoxir in OGC1 KD and scramble RNA transduced cells. Figure 3G shows that UK5099 increased etomoxir-sensitive OCR in cells transduced with scramble RNA, while in OGC1-KD cells this effect was abrogated. These data indicate that MPC inhibition requires glutamine metabolism and the MASh to allow increased fatty acid oxidation and energy expenditure.

**Figure 3:**
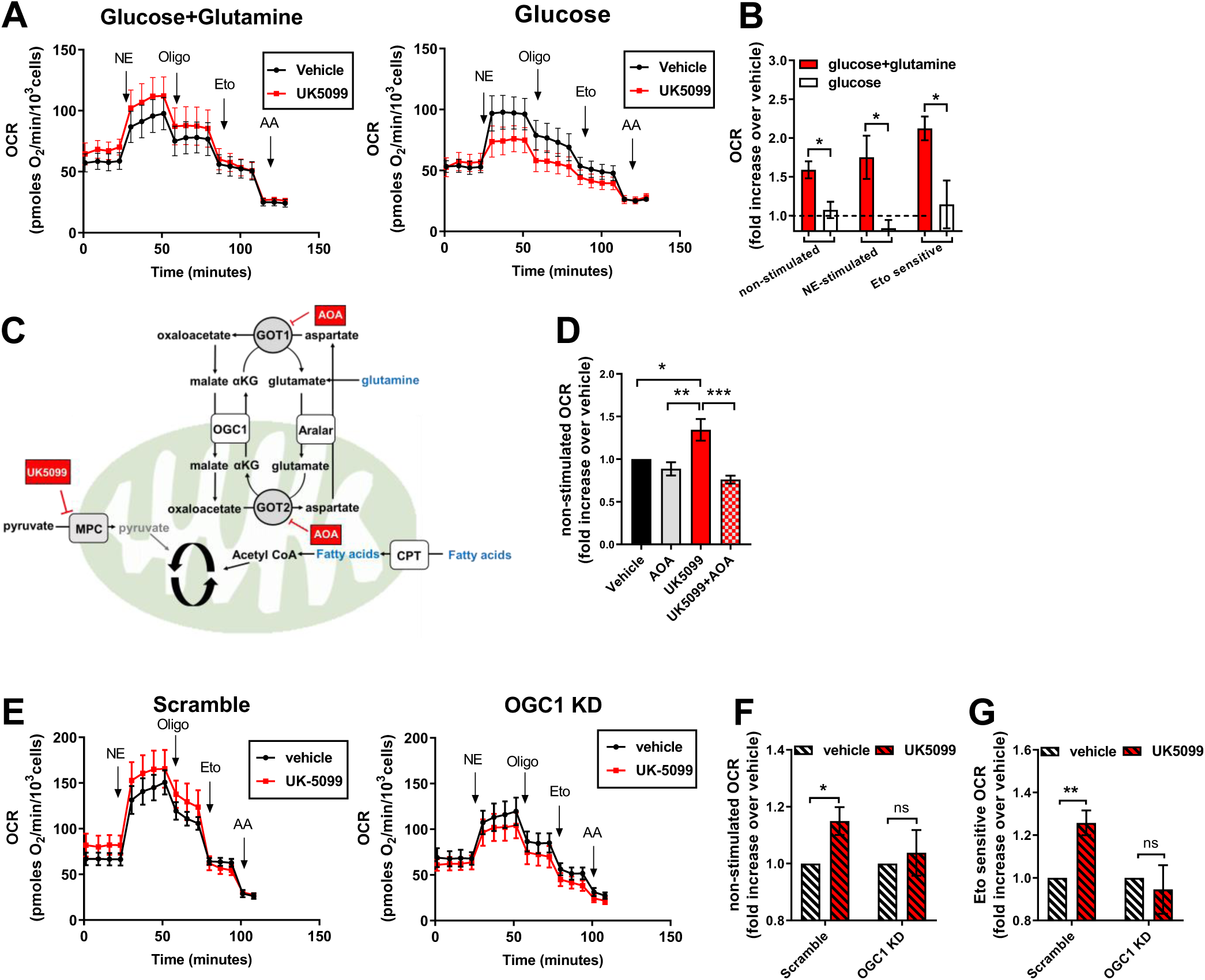
Increased fatty acid oxidation and energy expenditure promoted by MPC inhibition require glutamine metabolism and the activation of malate aspartate shuttle. **(A)** Primary brown adipocytes were pre-treated with vehicle (DMSO) or 100 nM UK5099 for 2 h. OCR was performed in presence of 5 mM glucose, or 5 mM glucose and 3 mM glutamine. 1 µM norepinephrine (NE), 4 µM oligomycin a (Oligo), 40 µM etomoxir (Eto) and 4 µM antimycin a (AA) were injected when indicated. Note that UK5099 increased OCR only when cells were assayed in the presence of both glucose and glutamine. Data shows representative OCR traces from n=6 technical replicates. **(B)** Quantification of basal and etomoxir-sensitive OCR after vehicle or UK5099 treatment (n=4 individual experiments). Data were normalized to vehicle for each experiment. ns p>0.05, ** p < 0.01, *** p < 0.001 compared to vehicle by Student’s t test. **(C)** Schematic of proposed mechanism of MPC inhibition induced activation of the malate-aspartate shuttle (MASh). GOT1/2, glutamic-oxaloacetic transaminase; OGC1, oxoglutarate carrier 1; Aralar, mitochondrial aspartate-glutamate carrier; CPT, Carnitine Palmitoyltransferase. **(D)** Brown adipocytes were pre-treated with vehicle (DMSO), 50-100 nM UK5099, 1 mM aminooxyacetic acid (AOA), or a combination of UK5099 and AOA for 2 h. Oxygen consumption rates (OCR) were measured in Seahorse base media supplemented with 5 mM glucose and 3 mM glutamine in the presence of the tested compounds. Data show quantification of basal OCR (n=5 individual experiments). Data were normalized to vehicle for each experiment. * p < 0.05, ** p < 0.01, *** p < 0.001 by ANOVA. **(E)** Primary brown adipocytes were transduced with Scramble or shOGC1 (OGC1 KD) adenovirus. Cells were pre-treated for 2 h with vehicle (DMSO) or 100 nM UK5099 before OCR measurements. OCR were measured in Seahorse base media supplemented with 5 mM glucose and 3 mM glutamine in the presence of vehicle or UK5099. 1 µM norepinephrine (NE, 4 µM oligomycin a (Oligo), 40 µM etomoxir (Eto) and 4 µM antimycin a (AA) were injected when indicated. Representative OCR traces with n=4 technical replicates. **(F)** Quantification of non-stimulated OCR (n=6 experiments). Data were normalized to vehicle for each experiment. * p < 0.05 compared to vehicle by Student’s t test. **(G)** Quantification of Eto-sensitive OCR (n=6 experiments). Data were normalized to vehicle for each experiment. * p < 0.05, ** p < 0.01 compared to vehicle by Student’s t test.

### 2.4. MPC inhibition increases ATP demand through activation of lipid cycling

We reasoned that increased OCR in non-stimulated brown adipocytes after MPC inhibition would result from either mitochondrial proton leak or increases in ATP demand. To assess the proportion of OCR that is linked to mitochondrial ATP synthesis, we used ATP synthase inhibitor oligomycin. The difference between OCR before and after oligomycin injection was calculated to determine ATP-linked OCR (Figure 4A). UK5099 increased oligomycin-sensitive OCR, suggesting that pharmacological inhibition of MPC increased mitochondrial ATP demand in brown adipocytes (Figures 4A-B). The contribution of ATP demand to the increase in energy expenditure mediated by MPC inhibition was seen at all UK5099 concentrations measured in a dose-response study (Figure S4A). ATP demand was also the main contributor to UK5099-mediated increase in energy expenditure in NE-stimulated cells, as revealed by OCR measurements of cells stimulated with NE prior to oligomycin injection (Figures 2A and 4B). We further observed an increase in mitochondrial proton leak in response to UK5099 treatment (Figure S4B), which could be a mechanism to maintain mitochondrial membrane potential. We next sought to unravel the mechanism behind the increase in ATP demand. We reasoned that activation of a futile cycle consuming ATP might explain the increased ATP demand observed upon MPC inhibition. Since MPC inhibition activated lipolysis and fatty acid oxidation (Figures 1C-E), we hypothesized that glycerolipid and free fatty acid cycling (GL/FFA, or lipid cycling) would represent the ATP demanding process activated by MPC inhibition (Figure 4C). Lipid cycling is a futile cycle of complete or partial lipolysis and lipid re-esterification, in which each complete cycle consumes 7 ATP molecules [27]. Indeed, lipid cycling represents an important component of basal energy demand and a key mechanism to prevent free fatty acid and glucose excess toxicity in various cell types [27], and plays an important role in adrenergically stimulated brown and beige adipocytes [28]. If lipid cycling is activated upon UK5099 treatment, we hypothesized that glycolytic intermediates should be increasingly diverted towards glycerol-3-phosphate (G3P) production. To test this possibility, we measured G3P and dihydroxyacetone-phosphate (DHAP) levels in brown adipocytes under UK5099 or NE treatment. Figure 4D shows that UK5099 increased the G3P/DHAP ratio similarly to NE, suggesting an increase in lipogenesis, thus supporting enhanced lipid cycling since this occurred in the face of accelerated lipolysis (Figure 1A). If lipid cycling is responsible for increased ATP demand upon UK5099 treatment, we reasoned that inhibition of lipolysis or lipid re-esterification should prevent UK5099-meditad increase in ATP demand. To test this hypothesis, we used pharmacological inhibitors of lipolysis or lipid re-esterification and assessed their effects on non-stimulated and ATP-linked OCR upon UK5099 treatment. Lipolysis was inhibited using Atglistatin, a specific inhibitor of adipose triglyceride lipase (ATGL), which mediates the first step in lipolysis. Figure 4E shows that Atglistatin completely reversed the UK5099-induced increase in non-stimulated OCR. Furthermore, Atglistatin fully reversed the effects of UK5099 on ATP-demand (Figure 4F). To address the possibility that inhibition of lipolysis on top of MPC inhibition prevented UK5099 mediated increase in OCR by affecting total respiratory capacity, we confirmed that treatment with Atglistatin and UK5099 did not affect FCCP stimulated OCR (Figure S4D). Lipid cycling can involve the entire cycle from TAG synthesis to free fatty acid and glycerol or be limited to shorter sub-cycles that involve break-down and regeneration of mono- and di-glycerides. Importantly, full- or partial cycling are both ATP-demanding processes. To assess the role of TAG synthesis step in UK5099-mediated activation of lipid cycling, we next determined the effect of diglyceride acyltransferase 1 and 2 (DGAT1/2) inhibitor. DGAT 1 and 2 catalyze the last step of TAG synthesis. We found that DGAT1/2 inhibition had no significant effect on non-stimulated or ATP-linked OCR in cells treated with UK5099 (Figure 4G and 4H). These results suggest that ATP-demand induced by MPC inhibition is not used for the last step of TAG synthesis. To assess whether UK5099 is activating partial lipid cycling, which does not require TAG synthesis, we used an inhibitor of Acyl-CoA synthetase (ACS), Triacsin C. We found that Triacsin C reverses the effects of UK5099 on non-stimulated and ATP-linked OCR back to vehicle OCR levels (Figure 4I and 4F). To assert that the inhibitory effect of Triacsin C on UK5099-induced OCR and ATP-demand is not a result of compromised respiratory capacity, we confirmed that treatment Triacsin C and UK5099 did not affect FCCP stimulated OCR (Figure S4E). Thus, our data suggest that UK5099 increases non-stimulated OCR and ATP-demand in brown adipocytes by activating a partial lipid cycling.

**Figure 4:**
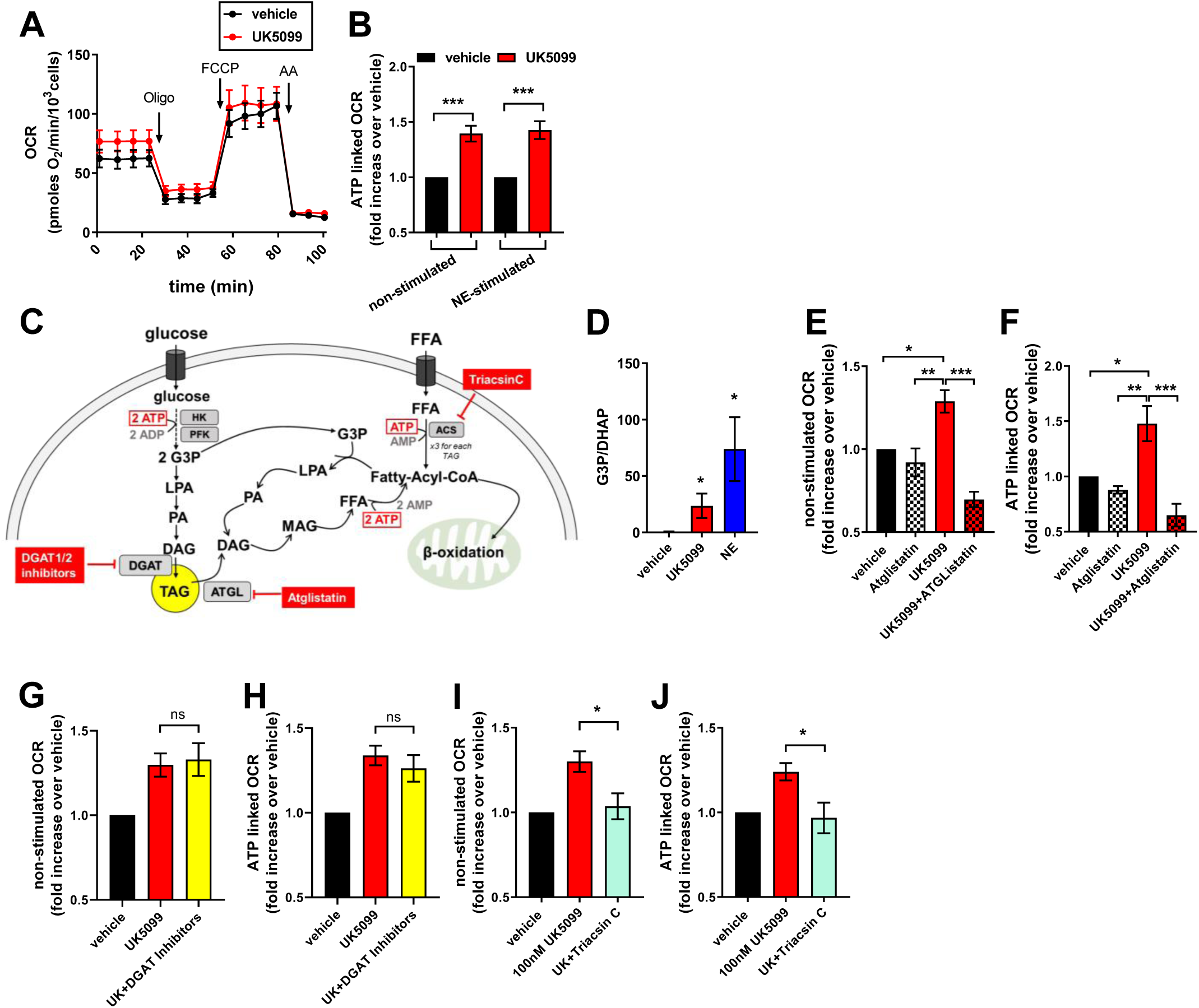
MPC inhibition increases ATP demand through activation of lipid cycling: **(A)** Primary brown adipocytes were pre-treated with vehicle (DMSO) or 100 nM UK5099 for 2 hours. OCR were measured in Seahorse base media supplemented with 5 mM glucose and 3 mM glutamine in the presence of vehicle or UK5099. 4 µM oligomycin a (Oligo), 2 µM mitochondrial uncoupler FCCP and 4 µM antimycin a (AA) were injected when indicated. Representative OCR traces with n=6 technical replicates. **(B)** Quantification of ATP-linked OCR (oligomycin sensitive) from n=9 experiments. ATP-linked respiration was calculated after injection at port A (basal), and after injection at port B following norepinephrine injection (NE). Data were normalized to vehicle for each individual experiment. *** p < 0.001 compared to vehicle by Student’s t test. **(C)** Schematic representation of lipid cycling. HK, hexokinase; PFK, phosphofructokinase; ATGL, adipose triglyceride lipase; ACS, Acyl-CoA synthetase; TAG, triacylglyceride; DAG, diacylglyceride; MAG, monoacylglyceride; LPA, lysophosphatidic acid; FFA, free fatty acid; G3P, glycerol-3-phosphate **(D)** Quantification of G3P to DHAP ratio in brown adipocytes treated with vehicle, 100 nM UK5099 or 1 µM norepinephrine (NE) for 24 h. Data shows n=4 individual experiments. * p < 0.05 by ANOVA. **(E)** Quantification of non-stimulated OCR from primary brown adipocytes treated with vehicle (DMSO), 100 nM UK5099, or 40 µM Atglistatin in combination with UK5099. Seahorse assay was performed as described in A. Data from n=4 experiments were normalized to vehicle for each individual experiment. * p < 0.05, ** p < 0.01, *** p < 0.001 by ANOVA. **(F)** Quantification of ATP-linked OCR from primary brown adipocytes treated with vehicle (DMSO), 100 nM UK5099, or 40 µM Atglistatin in combination with UK5099. Data from n=4 experiments were normalized to vehicle for each individual experiment. * p < 0.05, ** p < 0.01, *** p < 0.001 by ANOVA. **(G)** Quantification of non-stimulated OCR from primary brown adipocytes treated with vehicle (DMSO), 100 nM UK5099, or 1 µM DGAT1 and 1 µM DGAT2 inhibitors in combination with UK5099. Data from n=6 experiments were normalized to vehicle for each individual experiment. ns p>0.05 by ANOVA. **(H)** Quantification of ATP-linked OCR from primary brown adipocytes treated with vehicle (DMSO), 100 nM UK5099, or 1 µM DGAT1 and 1 µM DGAT2 inhibitors in combination with UK5099. Data from n=6 experiments were normalized to vehicle for each individual experiment. ns p>0.05 by ANOVA. (**I**) Quantification of non-stimulated OCR from primary brown adipocytes treated with vehicle (DMSO), 100 nM UK5099, or 5 µM Triacsin C in combination with UK5099. Data from n=4 experiments were normalized to vehicle for each individual experiment. * p < 0.05 by ANOVA. (**J**) Quantification of ATP-linked OCR from primary brown adipocytes treated with vehicle (DMSO), 100 nM UK5099, or 5 µM Triacsin C in combination with UK5099. Data from n=4 experiments were normalized to vehicle for each individual experiment. * p < 0.05 by ANOVA.

## 3. DISCUSSION

Our study identified a novel mechanism to activate futile lipid cycling and thereby increase brown adipocyte energy expenditure. Non-stimulated brown adipocytes store most of their intracellular lipids as lipid droplets, which are only consumed upon adrenergic stimulation [29]. Therefore, increasing fat oxidation in non-stimulated BAT, and potentially other types of adipose tissues, could be a promising way to reduce circulating fatty acids and shrink fat depots. Our data suggest that inhibition of the MPC increases lipolysis and fatty acid oxidation in both non-stimulated and NE-stimulated brown adipocytes. Fatty acid oxidation was assessed by measuring [U-^13^C_16_] palmitate incorporation into TCA cycle metabolites and using the CPT1 inhibitor etomoxir in respirometry assays. As etomoxir has been reported to have off-target effects particularly at higher doses [30], we confirmed that the dose of etomoxir used in this study does not affect the oxidation of substrates that are independent of transport via CPT1 (Figure S1). Interestingly, Vanderperre et al. showed that whole body MPC1 knock-out in mice was embryonically lethal, unless the pregnant mice were fed a ketogenic diet [16]. Further studies by Zou et al. showed that whole body heterozygous MPC1 knock-out mice have reduced body weight and increased expression of genes associated with lipolysis and fatty acid oxidation [31]. Additionally studies in C2C12 cells and muscle specific MPC1 KO mice have increased fatty acid oxidation [14,32]. This is in agreement with our results showing increased lipid droplet consumption and fatty acid oxidation in brown adipocytes treated with an MPC inhibitor. Our data also indicate that MPC inhibition acutely increases fatty acid oxidation and therefore could be a drug target for obesity and associated disorders via promoting fat oxidation and thermogenesis. Recent studies demonstrated that deleting MPC in Ucp1+ adipocytes from mice improved insulin sensitivity and increased circulating ketones, consistent with the increased fat oxidation detected in isolated brown adipocytes from these mice [33]. It remains to be determined whether MPC inhibition can promote fat oxidation and futile lipid cycling in white adipocytes, which would be especially interesting considering that most of human adipose tissue is white or beige.

Multiple studies have demonstrated that the modulation of MPC activity can act as a mitochondrial fuel preference switch in various tissues [14,17,31,34–37]. However, in these studies, pharmacological inhibition or genetic deletion of the MPC led to reduced or unchanged mitochondrial respiration. By contrast, we show that in brown adipocytes, inhibition of the MPC increased basal and adrenergically stimulated mitochondrial respiration. One possible reason that could explain this discrepancy is that the brown adipocyte is a cell type specialized to store and oxidize fat. Therefore, it is conceivable that brown adipocytes are more efficient for switching their fuel to fatty acids when mitochondrial pyruvate import is limited. Enhanced fatty acid oxidation is often associated with an increase in mitochondrial proton leak through UCP1 activation [38]. Surprisingly, we show that MPC inhibition indeed activated mitochondrial fatty acid oxidation but coupled to ATP synthesis by oxidative phosphorylation. Interestingly, we show that UK5099 increases ATP-linked OCR even in brown adipocytes stimulated with NE, suggesting that MPC inhibition can increase fat oxidation to produce ATP in mitochondria in which UCP1 was not successfully activated. In this regard, mitochondrial membrane potential measurements and IF staining of UCP1 show heterogeneity in these parameters between brown adipocytes, and between mitochondria inside the same cell [39].We show that MPC in brown adipocytes increases energy expenditure through the activation of ATP demanding lipid cycling. However, the mechanism by which inhibition of the MPC stimulates lipid cycling remains to be determined. It is possible that mitochondrial acetyl-CoA, PDH flux, the redox state or an intermediate of the TCA cycle regulates lipogenesis and lipolysis and thus free fatty availability towards oxidation. Nevertheless, our data support that glycolytic intermediates are diverted towards lipid cycling, instead of downstream glycolysis to pyruvate and lactate, as suggested by our data showing increased G3P/DHAP ratio upon MPC inhibition (Figure 4D). Further, because we observed that DAGT1/2 inhibition did not affect the action of MPC inhibition to promote fat oxidation, yet ATGL inhibition did, our data supports the view that lipogenesis up to TAGs, is not required for the acute action of the MPC inhibitor to promote TAG lipolysis, fat oxidation, lipid cycling and ATP usage, but supports the induction of a partial lipid cycling. As opposed to increasing energy wasting by mitochondrial uncoupling, activation of lipid cycling would be a safer and more controlled way to increase energy wasting. Indeed recent studies have proposed UCP1-independent futile cycles as possible mechanisms to increase energy expenditure in adipose tissue [7–10]. Future studies will determine whether lipid cycling can be activated by MPC inhibition in white and beige adipose tissue as well.

In addition to increased oxidation of fatty acids, our data indicate that glutamine supports energy expenditure induced by MPC inhibition, as suggested by previous studies [14,17,37]. Intriguingly, we find that MPC inhibition in brown adipocytes requires the MASh activity to increase oxygen consumption rates. In this sense, glutamine and the MASh provide two mechanism to increase respiratory rates: i) by allowing its condensation with acetyl-CoA from beta-oxidation to maintain TCA cycle and ii) by providing a mechanism for cytosolic NADH to be used by electron transport chain in mitochondria through the MASh activity. Therefore, the identification that MASh enhances energy expenditure in brown adipocytes (or MAShEEBA) represents a novel mechanism for regulation of energy metabolism in BAT. Although the mechanism by which the MASh is activated upon MPC remains to be determined, it is conceivable that it is directly linked to the activation of lipolysis and increased fatty acid oxidation upon MPC inhibition. Indeed, Wang et al. showed that acetyl-CoA derived from fatty acids can acetylate and thereby activate components of the MASh [40]. Interestingly the only described posttranslational modification of the MPC is acetylation of MPC2, which was shown to have an inhibitory effect on MPC activity [41]. The observation that acetylation has activating effects on the MASh, and inhibitory effects on the MPC might suggest a functional link between MASh activity and MPC activity. The role of the MASh under physiological stimulation of non-shivering thermogenesis remains to be determined.

In conclusion, we identified a novel mechanism to activate futile lipid cycling and increase energy expenditure through the inhibition of the MPC. Importantly, FDA-approved drugs were shown to be target the MPC at clinically relevant concentration [42]. This suggests that the MPC could be a safe target to increase energy expenditure in brown adipocytes and potentially improve whole body metabolic health in patients with obesity and cardiometabolic disorders linked to nutrient excess.

## 4. METHODS

### 4.1. Animals

Primary brown adipocytes were isolated from 4-5 weeks old wild-type male C57BL/6J mice (Jackson lab, Bar Harbor, ME). Animals were fed standard chow (mouse diet 9F, PMI Nutrition International, Brentwood, MO) and maintained under controlled conditions (19–22°C and a 14:10 hour light-dark cycle) until euthanasia by isofluorane, followed by cervical dislocation. All animal procedures were performed in accordance with the Guide for Care and Use of Laboratory Animals of the NIH and were approved by the ARC/IACUC of the University of California, Los Angeles.

### 4.2. Primary brown adipocyte culture

Primary brown adipocytes were isolated and cultured as described in detail [1]. In brief, BAT was dissected from interscapular, subscapular, and cervical regions of four male mice. Tissue was digested using Collagenase Type II (Worthington, Lakewood, NJ). Digested tissue was filtered through a 100 µm and 40 µm mesh and centrifuged. Cell pellet was re-suspended in brown adipocyte culture media (DMEM supplemented with 10 % newborn calf serum (Sigma-Aldrich, St. Louis, MO), 4 mM glutamine, 10 mM HEPES, 0.1 mg/mL sodium ascorbate, 100 U/mL penicillin, 100 µ/mL streptomycin) and plated in a 6-well plate. Cells were incubated in a 37°C, 8 % CO_2_ incubator. 72 hours after isolation the cells were lifted using STEMPro Accutase (Thermo Fisher Scientific, Roskilde, Denmark), counted, and re-plated in final experimental vessel. 24 hours later media was changed to differentiation media (growth media supplemented with 1 µM rosiglitazone maleate (Sigma-Aldrich, St. Louis, MO) and 4 nM human insulin (Humulin R, Eli Lilly, Indianapolis, IN). Cells were differentiated for 7 days and media was changed every other day.

### 4.3. Gene silencing

#### 4.3.1. Adenoviral Transduction

On day 3 of differentiation, adipocytes were incubated with 1.5 µL/mL of adenoviral preparation (10^9^ particles/ml) for 24 h in complete culture media containing 1 µg/mL polybrene (hexadimethrine bromide, Sigma-Aldrich, St. Louis, MO). Respirometry, metabolomics and gene expression were measured on day 7 of differentiation. Ad-mSLC25A11 and Ad-mKate2 shControl were generated and purchased from Welgen (Worcester, MA).

#### 4.3.2. siRNA transfection

Undifferentiated pre-adipocytes were transfected with scramble RNA or MPC1 siRNA using Lipofectamine 3000 reagent (Thermo Fisher Scientific, Roskilde, Denmark) according to manufacturer’s protocol. In brief; culture media was removed from cells and cells were incubated with Opti-MEM media (Thermo Fisher Scientific, Roskilde, Denmark), Lipofectamine 3000 reagent and scramble RNA or siRNA for 4 hours. Then DMEM with 1 % fetal bovine serum (Thermo Fisher Scientific, Roskilde, Denmark) was added to the cells and incubated overnight. The next day media was replaced with differentiation media. Respirometry, gene expression and protein expression were measured on day 7 of differentiation. The following siRNAs were used: ON-TARGETplus Mouse Mpc1 (55951) siRNA (L-040908-01-0005) and ON-TARGETplus Non-targeting Pool (D-001810-10-05) from Dharmacon (Lafayette, CO).

### 4.4. Respirometry measurements

#### Respirometry in intact cells

Pre-treatments were performed in brown adipocyte culture media. The following compounds were used for pre-treatments: 50 nM - 20 µM UK5099 (Sigma-Aldrich, St. Louis, MO), 40 µM Atglistatin (Selleck Chemicals, Houston, TX), 200 µM – 1 mM aminooxyacetic acid (Sigma-Aldrich, St. Louis, MO). Prior to respirometry measurements, culture media was replaced to Seahorse assay media (Seahorse XF Base medium (Agilent Technologies, Santa Clara, CA) supplemented with 5 mM glucose and 3 mM glutamine and incubated for 30-45 minutes at 37°C (without CO_2_). When indicated, seahorse media was supplemented with 0.1 % fatty acid free bovine serum albumin (Sigma-Aldrich, St. Louis, MO), or 5 % fetal bovine serum (Thermo Fisher Scientific, Roskilde, Denmark). During this incubation period, the ports of the Seahorse cartridge were loaded with the compounds to be injected during the assay (50 μL/port) and the cartridge was calibrated. Oxygen consumption rates were measured using the Seahorse XF24-3 extracellular flux analyzer (Agilent Technologies, Santa Clara, CA). The following compounds were used for injections during the assay: 1 µM norepinephrine, 4 µM oligomycin a (Calbiochem, San Diego, CA), 40 µM etomoxir (Sigma-Aldrich, St. Louis, MO), 4 µM antimycin a (Sigma-Aldrich, St. Louis, MO), 2 µM FCCP (Sigma-Aldrich, St. Louis, MO). After the assay, cells were fixed using 4 % paraformaldehyde (Thermo Fisher Scientific, Roskilde, Denmark). To normalize the data for possible differences in cell number, nuclei were stained with 1 µg/mL Hoechst 33342 (Thermo Fisher Scientific, Roskilde, Denmark) and nuclei were counted using the Operetta High-Content Imaging System (PerkinElmer, Waltham, MA).

#### Respirometry in permeabilized cells

Experiments were performed as previously described in detail [43]. In brief, differentiated primary brown adipocytes were permeabilized using 7.5 nM XF PMP reagent (Agilent Technologies, Santa Clara, CA). Respirometry assay was performed in MAS buffer (660 mM mannitol, 210 mM sucrose, 30 mM KH_2_PO_4_, MgCl_2_, HEPES, EGTA, 1% (w/v) fatty-acid free BSA). The following substrates were used: 100 μM palmitoyl-carnitine+3 mM malate+500 μM carnitine+4 mM ADP, or 100 μM palmitoyl-CoA+3 mM malate+ 500 μM carnitine+ 4 mM ADP. Oligomycin was injected at 5 μM and antimycin a at 8 μM final concentration. Oxygen consumption rates were measured using the Seahorse XF24-3 extracellular flux analyzer (Agilent Technologies, Santa Clara, CA).

### 4.5. Quantitative real time PCR

Samples from cells transfected with shRNA (OGC1-KD) or siRNA (MPC1-KD) were collected on day 7 of differentiation. RNA was extracted using the Direct-zol RNA Miniprep Plus Kit ® (Zymo Research, Irvine, CA) following the manufacturer’s instructions. A sample corresponding to 1 µg RNA from each sample was used to perform cDNA synthesis by the High-Capacity cDNA Reverse Transcription Kit ® (Applied Biosystems, Foster City, CA). QPCR were performed using 0.4 ng/µL cDNA and 240 nM of each primer, whose sequences are listed in Table I.

**Table 1:**
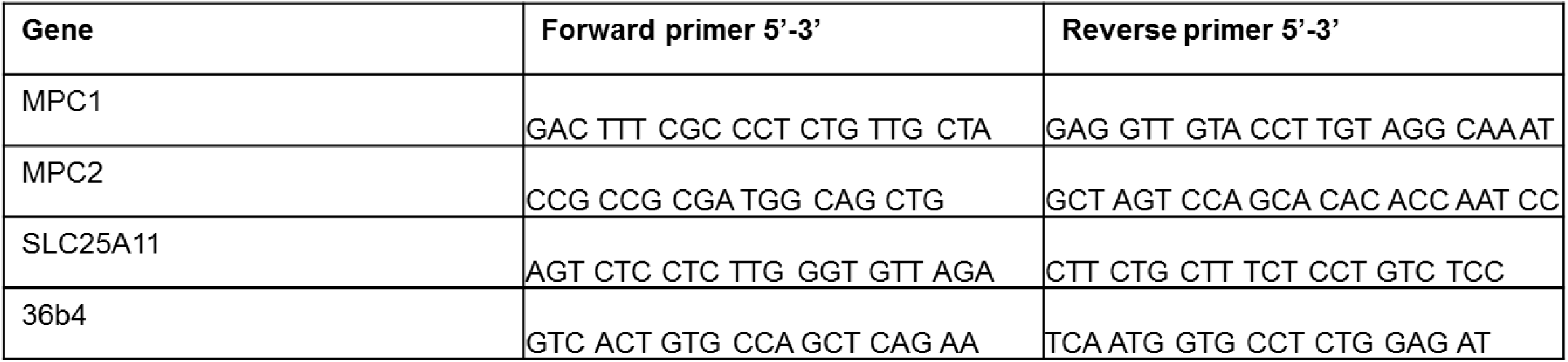
Primers for qPCR.

### 4.6. Western blot

Protein was isolated using RIPA lysis buffer (Santa Cruz Biotechnology, Dallas, TX) containing protease and phosphatase inhibitor cocktail (Thermo Fisher Scientific, Roskilde, Denmark), and protein concentration was determined using a BCA protein assay (Thermo Fisher Scientific, Roskilde, Denmark). Samples of 10-15 µg isolated protein were diluted in NuPAGE LDS Sample Buffer (Thermo Fisher Scientific, Roskilde, Denmark) containing β-mercaptoethanol (Thermo Fisher Scientific, Roskilde, Denmark), and incubated at 95°C for 5 min. Samples were then loaded into 4 %– 12 % Bis-Tris precast gels (Thermo Fisher Scientific, Roskilde, Denmark) and electrophoresed, using NuPAGE MES SDS Running Buffer (Thermo Fisher Scientific, Roskilde, Denmark). Proteins were transferred to methanol-activated Immuno-Blot PVDF Membrane (Bio-Rad, Hercules, CA). Blots were incubated over-night with primary antibody diluted in PBST (phosphate buffered saline with 1 mL/L Tween-20/PBS) + 5 % BSA (Thermo Fisher Scientific, Roskilde, Denmark) at 4°C. The next day, blots were washed in PBST and incubated with fluorescent secondary antibodies, diluted in PBST+ 5 % BSA for 1 hour at room temperature. Proteins were detected using the following antibodies: anti-MPC1 (BRP44L Polyclonal Antibody, Thermo Fisher Scientific, Roskilde, Denmark), anti-MPC2 (MPC2 (D4I7G) Rabbit mAb, Cell Signaling, Danvers, MA), anti-β-Actin (Abcam, Cambridge, United Kingdom), Goat anti-Rabbit IgG secondary antibody DyLight 800 (Thermo Fisher Scientific, Roskilde, Denmark). Blots were imaged on the ChemiDoc MP imaging systmen (Bio-Rad Laboratories, Hercules, CA). Band densitometry was quantified using FIJI (ImageJ, NIH).

### 4.7. Fluorescence confocal microscopy

Super resolution live cell imaging was performed on a Zeiss LSM880 using a 63x Plan-Apochromat oil-immersion lens and AiryScan super-resolution detector with humidified 5 % CO_2_ chamber on a temperature controlled stage (37°C). Cells were differentiated in glass-bottom confocal plates (Greiner Bio-One, Kremsmünster, Austria). One day prior to imaging, cells were incubated with 1 µM Bodipy 558/568 C12 (Thermo Fisher Scientific, Roskilde, Denmark) overnight. The day of imaging cells were stained with 200 nM mitotracker green (Thermo Fisher Scientific, Roskilde, Denmark) for 1 h. mitotracker green and Bodipy 558/568 C12 were removed before imaging and cells were imaged in regular culture media. Mitotracker green was excited with 488 nm laser and Bodipy 558/568 C12 was excited with 561 nm laser. Image Analysis was performed in FIJI (ImageJ, NIH). Image contrast and brightness were not altered in any quantitative image analysis protocols. Brightness and contrast were equivalently modified in the different groups compared, to allow proper representative visualization of the effects revealed by unbiased quantitation.

### 4.8. Thin layer chromatography

Cells were seeded and differentiated in 6-well plate. On day 7 of differentiation, cells were incubated with 1 µM Bodipy 558/568 C12 and DMSO, 100 nM UK5099 or 1 µM norepinephrine for 4 h and 24 h. Thin layer chromatography of intra-cellular lipids and extra-cellular lipids was performed as previously described [20] with minor modifications. Intra-cellular lipids and lipids from media were extracted in 500 µL chloroform. Chloroform was evaporated using the Genevac EZ-2 Plus Evaporating System (Genevac, Ipswich, United Kingdom). Lipids were then dissolved in 15 µL Chloroform and 1 µL was spotted on a TLC plate (Silica gel on TLC Al foils, Sigma-Aldrich, St. Louis, MO). Lipids were resolved based on polarity in a developer solution containing ethylacetate and cyclohexane in a 2.5:1 ratio. TLC plates were imaged on the ChemiDoc MP imaging systmen (Bio-Rad Laboratories, Hercules, CA). Band densitometry was quantified using FIJI (ImageJ, NIH).

### 4.9. Metabolite tracing

For metabolite, tracing cells were cultured and differentiated in 6-well plates using BAT differentiation media. On day 7 of differentiation cells were washed once and treated with 5 µM UK5099 or DMSO for 24 h in DMEM supplemented with phenol red, 10 mM glucose, 2 mM glutamine, 200 µM [U-^13^C_16_] palmitate (Cambridge Isotope Laboratories, Tewksbury, MA) and 10% delipidated NCS. Palmitate was added at a 4:1 palmitate:BSA complex. NCS was delipidated using fumed silica (Sigma-Aldrich, St. Louis, MO). Metabolite extraction and GC/MS was performed as previously described in detail [14].

### 4.10. G3P and DHAP measurements

G3P and DHAP were measured using fluorimetric and colorimetric kits available from Sigma-Aldrich (MAK207 and MAK275, respectively). After differentiation in 6-well plates, cells were treated with DMSO (vehicle), 100 nM UK5099 or 1 µM NE for 24 hours. Then, cells were washed with PBS and 60 µL of the respective assay buffer were added to each well. The suspension was centrifuged for 2,000 x g for 1 min and 10 µL of supernatant were used to measure G3P and DHAP according to the manufacturer instructions. For normalization purposes, 2 µL were used to measure protein concentration by a BCA protein assay kit (Thermo Fisher Scientific, Roskilde, Denmark).

### 4.11. Statistical analyses

Data were presented as mean ± SEM for all conditions. D’Agostino and Pearson normality test was performed to assess Gaussian distribution. Comparisons between groups were done by one-way ANOVA and *a posteriori* Tukey’s test for pair-wise comparisons. When appropriate, two-way ANOVA, unpaired/paired Student’s t-tests or Mann-Whitney’s test were employed. Differences of p<0.05 were considered to be significant. All graphs and statistical analyses were performed using GraphPad Prism 7 for Windows (GraphPad Software, San Diego, CA).

## ACKNOWLEDGEMENTS

OSS is funded by NIH-NIDDK 5-RO1DK099618-02. MFO is funded by Conselho Nacional de Desenvolvimento Científico e Tecnológico (grant # 229526/2013-6). ML is funded by the Department of Medicine chair commitment at UCLA, Pilot and Feasibility grants from NCATS UL1TR001881 (CTSI), NIDDK P30 DK063491 (UCSD-UCLA DERC) and NIDDK P30 41301 (CURE:Digestive Diseases Research Center). AEJ is funded by the UCLA Tumor Cell Biology Training Program (USHHS Ruth L. Kirschstein Institutional National Research Service Award #T32 CA009056). Special thanks to Barbara Cannon and Jan Nedergaard for guiding our labs into the field of thermogenesis and brown adipocyte biology.

## AUTHOR CONTRIBUTIONS

Conceptualization; M.V., M.L., M.F.O., O.S.S, Investigation; M.V., C.M.F., I.Y.B., A.E.J., B.R.D., K.M., R.A-P., A.P., Writing-Original Draft; M.V., M.F.O., M.L., O.S.S. Writing-Review & Editing; M.V., A.S.D., M.P., B.E.C., M.L., M.F.O., O.S.S

## Figure legends

**Figure S1.**
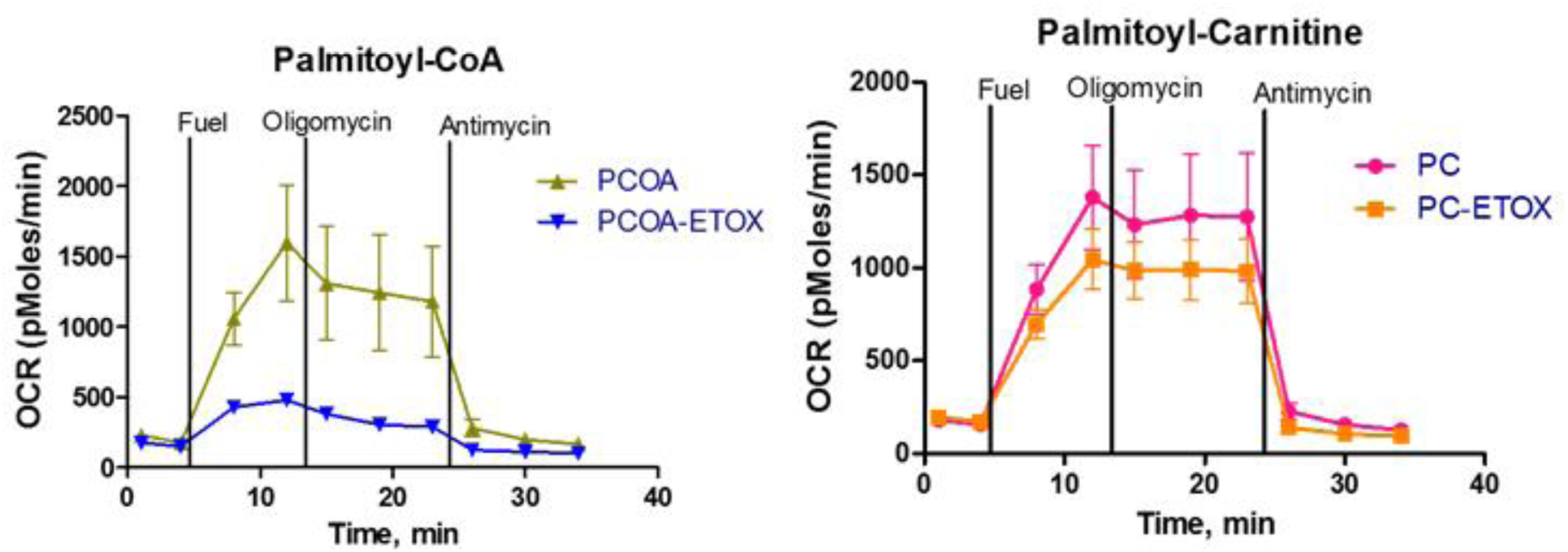
OCR traces of permeabilized primary brown adipocytes. OCR were measures in the presence of palmitoyl-CoA, a substrate that is dependent on CPT1, or in the presence of palmitoyl-carnitine a substrate that does not require CPT1 activity. Note that etomoxir only inhibits OCR fueled by palmitoyl-CoA.

**Figure S2:**
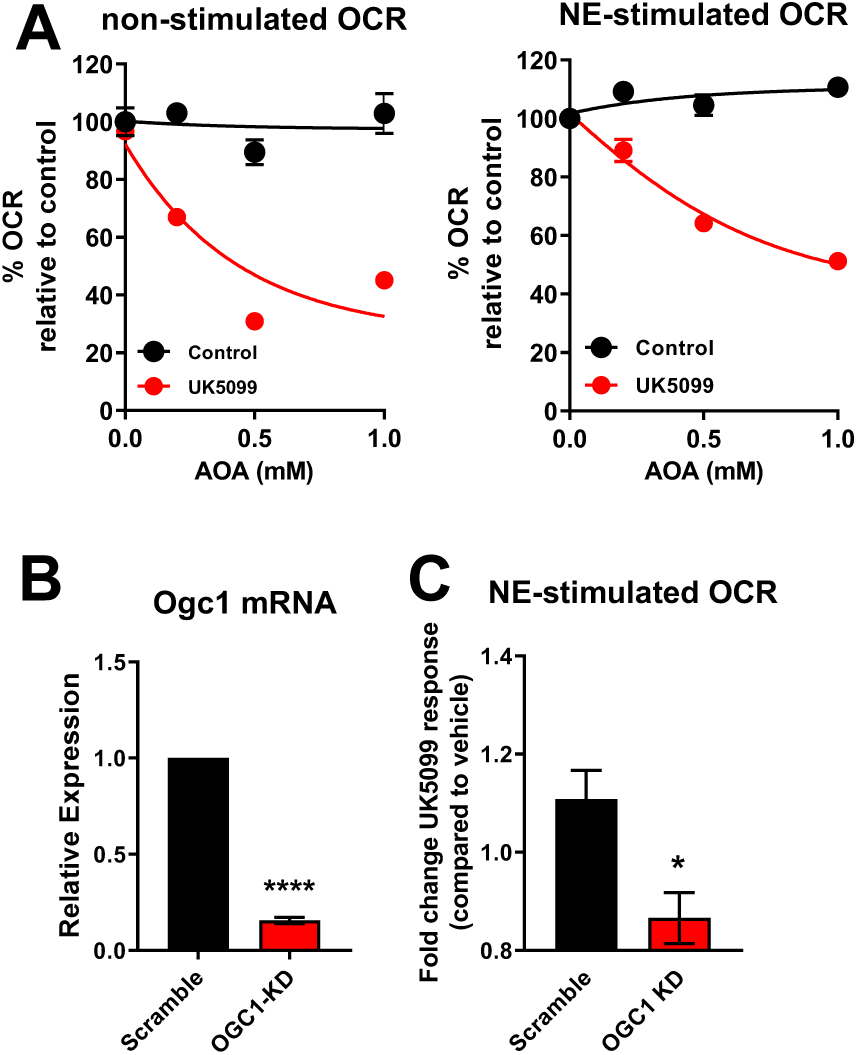
**(A)** Brown adipocytes were pre-treated with vehicle (DMSO), UK5099 (50 nM), aminooxyacetic acid (AOA) at various concentrations, or a combination. Oxygen consumption rates (OCR) were measured in Seahorse base media supplemented with 3 mM glucose and 3 mM glutamine in the presence of the compounds. Data shows non-stimulated and norepinephrine stimulated OCR. Note that AOA has no effect in vehicle treated cells, but reduced OCR when cells are treated with UK5099. **(B)** mRNA levels of OGC1 in brown adipocytes transduced with adenovirus carrying either scramble RNA (Scramble) or OGC1 shRNA (OGC1-KD). **** p < 0.0001 compared to Scramble by Student’s t-test. **(C)** Quantification of norepinephrine stimulated OCR after vehicle or UK5099 treatment in scramble RNA or OGC1 shRNA transduced cells. Data were normalized to vehicle for each individual experiment. * p < 0.05 compared to vehicle by Student’s t-test.

**Figure S4:**
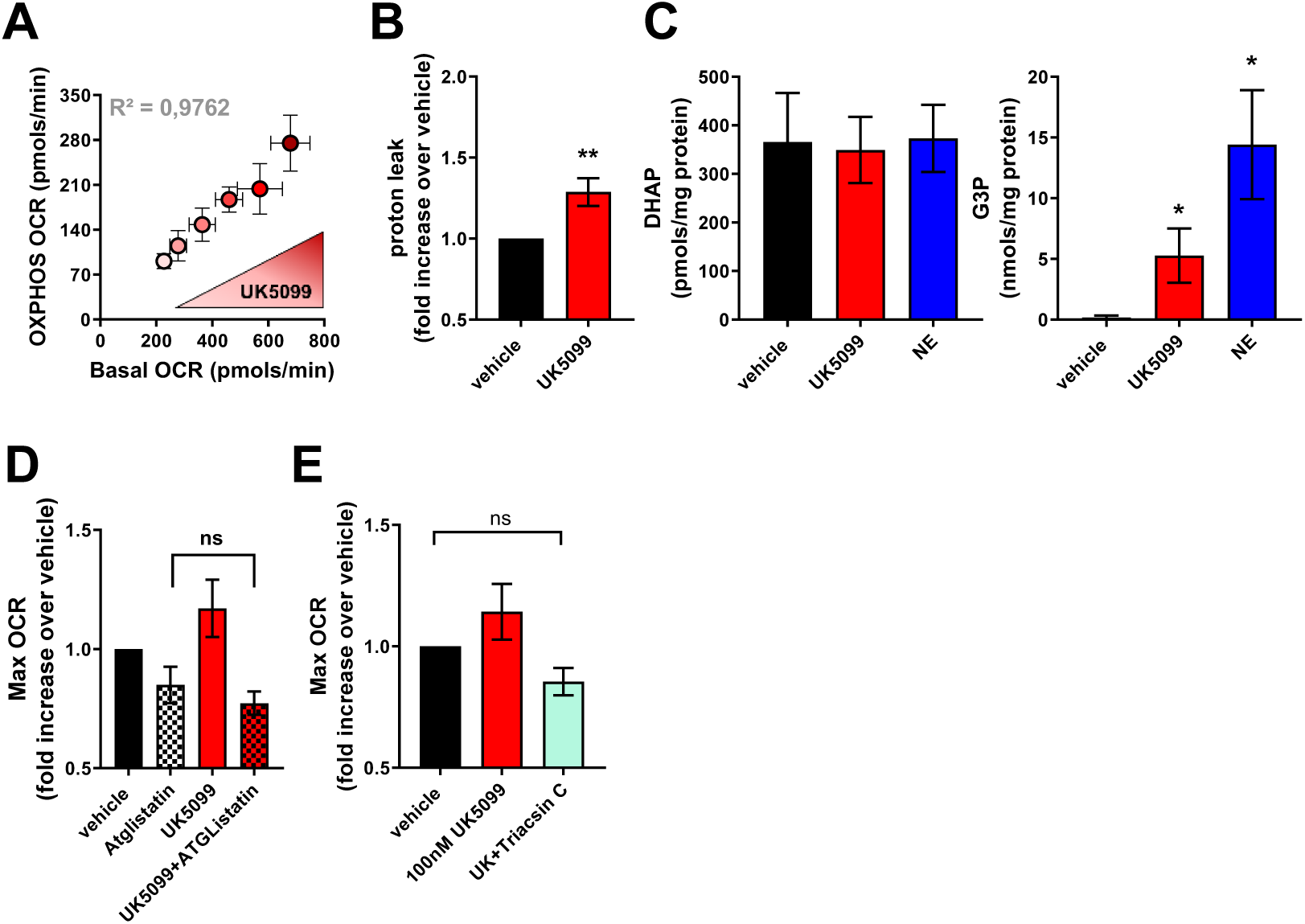
**(A)** Brown adipocytes were treated with increasing concentration of UK5099. Data shows changes in basal OCR as a function of ATP linked OCR. Note that the dose dependent increase in basal OCR following UK5099 treatment correlates with an increase in ATP linked OCR. **(B)** Quantification of oligomycin insensitive proton leak in brown adipocytes treated with 100 nM UK5099 or vehicle (n=10 individual experiments). Data were normalized to vehicle for each individual experiment. ** p < 0.01 compared to vehicle by Student’s t test. **(C)** Quantification of G3P and DHAP in brown adipocytes treated with vehicle, 100 nM UK5099 or 1 µM norepinephrine (NE) for 24 h. * p < 0.05 compared to vehicle by ANOVA. **(D)** Brown adipocytes were treated with vehicle (DMSO), 100 nM UK5099 or 40 µM Atglistatin in combination with UK5099. Maximal OCR (Max OCR) were calculated as maximal OCR after FCCP injection. Data were normalized to vehicle for each individual experiment. ns p>0.05 compared to vehicle by ANOVA. (**E**) Brown adipocytes were treated with vehicle (DMSO), 100 nM UK5099 or 5 µM Triacsin C in combination with UK5099. Maximal OCR (Max OCR) were calculated as maximal OCR after FCCP injection. Data were normalized to vehicle for each individual experiment. ns p>0.05 compared to vehicle by ANOVA.

